# One million years of solitude: the rapid evolution of de novo protein structure and complex

**DOI:** 10.1101/2023.12.24.573215

**Authors:** Jianhai Chen, Qingrong Li, Shengqian Xia, Deanna Arsala, Dylan Sosa, Dong Wang, Manyuan Long

## Abstract

Recent studies have established that de novo genes, evolving from non-coding sequences, enhance protein diversity through a stepwise process. However, the pattern and rate of their structural evolution over time remain unclear. Here, we addressed these issues within a short evolutionary timeframe (∼1 million years for 97% of rice de novo genes). We found that de novo genes evolve faster than gene duplicates in the intrinsic disordered regions (IDRs, such as random coils), secondary structural elements (such as α-helix and β-strand), hydrophobicity, and molecular recognition features (MoRFs). Specifically, we observed an 8-14% decay in random coils and IDR lengths per million years per protein, and a 2.3-6.5% increase in structured elements, hydrophobicity, and MoRFs. These patterns of structural evolution align with changes in amino acid composition over time. We also revealed significantly higher positive charges but smaller molecular weights for de novo proteins than duplicates. Tertiary structure predictions demonstrated that most de novo proteins, though not typically well-folded on their own, readily form low-energy and compact complexes with extensive residue contacts and conformational flexibility, suggesting “a faster-binding” scenario in de novo proteins to promote interaction. Our findings illuminate the rapid evolution of protein structure in the early life of de novo proteins in rice genome, originating from noncoding sequences, highlighting their quick transformation into active, complex-forming components within a remarkably short evolutionary timeframe.

## Introduction

The complexity and adaptability of biological systems often find their roots in the ever-budding genetic landscape. Central to this is the emergence of de novo genes -- genes that arise from regions of DNA once categorized as ’junk’ and considered functionally insignificant (Fagundes, et al. 2022; Ohno 1972). The birth of de novo genes was deemed impossible or functionally irrelevant (Jacob 1977; Mayr 1982). However, recent studies have challenged this dogma and provided concrete evidence that de novo genes can indeed emerge from non-coding sequences through a stepwise mutational process, contributing to increased protein diversity (Heames, et al. 2020; Zhang, et al. 2019). Despite these progresses, our understanding of these novel proteins, particularly their structural characteristics at the secondary, tertiary, and complex levels, and the rate of their structural evolution, remains largely unexplored.

Gene duplicates have long been recognized as a predominant source of new genes. These duplicates retain sequences from their parent genes and contribute to phenotypic evolution through various mechanisms, including neofunctionalization, hypofunctionalization, subfunctionalization and gene dosage (Birchler and Yang 2022; Kaessmann 2010; Ohno 1970). In contrast, de novo genes evolve through non-duplication mechanisms and have been shown to play diverse roles in biological functions. Their contributions have been highlighted in multiple systems, including DNA repair in yeast (Cai, et al. 2008), providing a novel antifreeze function in Arctic fish (Zhuang and Cheng 2021), diversification of rice morphology (Chen, et al. 2023b), cortical expansion in humans (An, et al. 2023; Qi, et al. 2023), and even oncogenesis in human cancers (Suenaga, et al. 2014). The emergence and functional diversity of de novo genes introduce a novel dimension to our understanding of genome evolution and functional innovation, expanding our knowledge beyond traditional gene duplication models (Broeils, et al. 2023; Carvunis, et al. 2012; Knowles and McLysaght 2009; Vakirlis, et al. 2022; Zhang, et al. 2019; Zhao, et al. 2014).

Due to their relatively recent origin, de novo proteins may not have evolved into well-folded structure. This leads to a characteristic feature: a lack of stable tertiary structure when isolated, thus manifesting as intrinsically structural disorder (ISD) and extensive formation of intrinsic disordered regions (IDRs) or random coils. It is found that exquisitely adapted species contains more ISD domains (Weibel, et al. 2023). ISD are also commonly found in proteins related to human genetic diseases (Midic, et al. 2009; Vavouri, et al. 2009). Despite advancements in function studies of ISD proteins, the extent of ISD in de novo genes remains a subject of debate. Several studies suggest a strong tendency towards ISD in de novo genes or newly evolved domains (Basile, et al. 2017; Bitard-Feildel, et al. 2015; Heames, et al. 2023; Heames, et al. 2020; Lange, et al. 2021; Wilson, et al. 2017). Conversely, other studies present conflicting results, contradicting the association between gene sequence novelty and ISD or suggesting no apparent correlation (Ekman and Elofsson 2010; Schmitz, et al. 2018; Vakirlis, et al. 2018). Furthermore, the question of whether ISD is influenced by gene age or if it can evolve over time remains unresolved.

Additionally, the evolvability of well-folded structural elements in de novo genes, such as, 3_10_ helices, α-helices, and β-strands etc., remains an open question. If de novo proteins can gradually evolve from a disordered to a well-folded structure, an intriguing question arises: how are their sequence compositions optimized for structural stability over evolutionary time? With advances like AlphaFold heralding a new era in protein structure prediction (Jumper, et al. 2021), we can now conduct an in-depth exploration of the evolution of de novo protein structure and elements over evolutionary time. Other questions also include how could these de novo proteins, which are often very short, interact with other usually larger proteins, and their ability to form complexes with other biomolecules. Indeed, roughly 40% of all protein-protein interactions are between proteins and shorter peptides, many of which play critical roles in cellular life-cycle functions (Lee, et al. 2019). Recent advances like AlphaFold-Multimer excels in predicting peptide-protein interactions (Johansson-Åkhe and Wallner 2022), which could facilitate our understanding on the evolution of de novo protein and potential conformational changes upon binding.

The structural evolution of proteins is conventionally perceived as a slow process, maintaining remarkable conservation over hundreds of millions to billions of years, in contrasts with the rapid changes observed in their primary structure (Ingles-Prieto, et al. 2013; Liljas, et al. 2016). In this conventional view, within the relatively short evolutionary timescale of one to a few million years, it was assumed that little to no significant structural changes would occur in proteins, let alone the emergence of new protein-protein interactions. Thus, the evolution of protein structure could be perceived as a process of “million years of solitude.”

In this study, we explore the evolutionary patterns of de novo genes with a focus on their protein structures and complexes. We analyzed multiple properties of protein structure including the proportions of intrinsic disordered regions (IDRs), secondary structure elements (including the unstructured random coils and structured α-helices and β-strands), amino acid composition and properties (such as charges, weights, and hydrophobicity), molecular recognition features (MoRFs), and the protein complexes. We revealed the rapid evolution of de novo proteins in forming structures and complexes due to their different features from duplicated proteins, which could reshape our understanding on new gene evolution. These insights challenge the conventional view of protein evolution and have revealed a dynamic world of protein evolution over short evolutionary timescales.

## Results

### The levels of ISD in de novo proteins reduce gradually over evolutionary time

We retrieved gene list of de novo genes from our previous study, which demonstrated detailed stepwise process of de novo gene emerge from non-coding regions (Zhang, et al. 2019) (Figure 1a). The gene ages are defined as the branches with the complete Open Reading Frame (ORF) formation. We locally inferred gene ages based on the synteny of reciprocal best neighbor genes for 27,673 duplicated genes (Long, et al. 2013), which account for 71.41% of genomic protein-coding genes (IRGSP-1.0.75 version of rice genome) (Figure 1b). Both gene duplicates and *de novo* genes were assigned into evolutionary age groups from young to old evolutionary epochs based on their known phylogenetic framework (Zhang, et al. 2019) (Figure 1c, Supplementary table 1). The nine evolutionary age groups cover ∼15 million years of Oryzeae evolution, which includes species of *O. sativa japonica* (br1), *O. rufipogon* (br2), *O. sativa* subspecies *indica* and *O. nivara* (br3), *O. glaberrima* and *O. barthii* (br4), *O. glumaepatula* (br5), *O. meridionalis* (br6), *O. punctata* (br7), *O. brachyantha* (br8), and *L. perrieri* (br9) (Figure 1c). We found 97% of rice de novo genes are within the timeframe of around one million years (br1-br5, 169/175).

**Figure 1.**
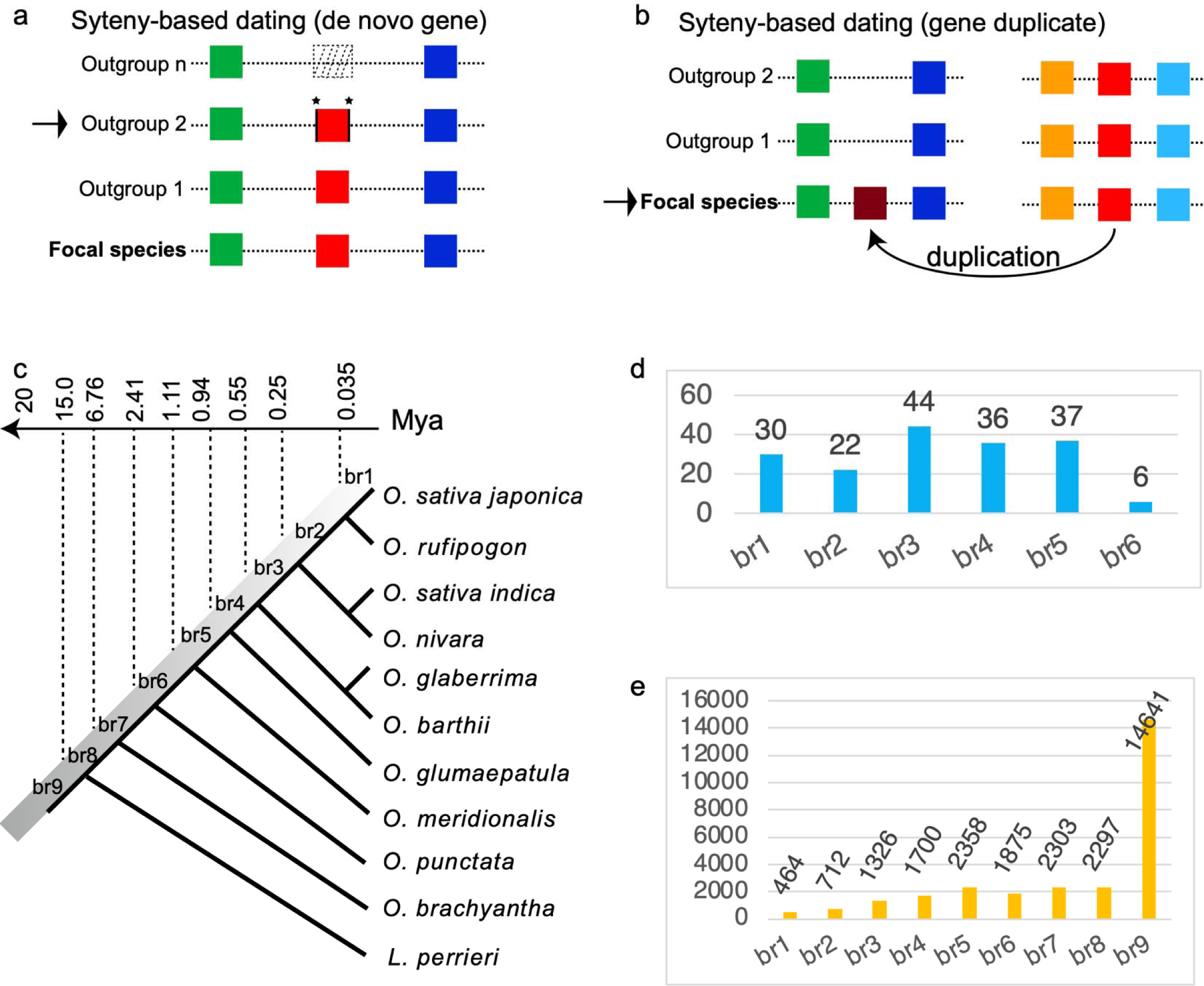
The methodology of gene age dating and number of genes with gene age information for de novo genes and gene duplicates. (a) The method for dating de novo gene ages, based on our previous synteny-based study (Zhang, et al. 2019). The dotted box indicates non-coding sequence with DNA-level similarity to de novo genes (red). The neighboring genes are represented in green and blue, with outgroup 2 containing a complete ORF. The emergence of the gene is attributed to ’trigger’ or ’enabler’ mutations, including substitutions and/or insertions/deletions (indicated by asterisks), as detailed in (Zhang, et al. 2019). (b) For duplicated genes, the synteny-based age dating method involves identifying homologous genes (depicted in purple and red) and their neighboring genes. The direction of duplication is indicated by an arrow. The emergence of the purple gene is determined based on the presence or absence of conserved synteny in the focal species. (c) The phylogenetic framework (br1-br9) and the corresponding divergence time (million years ago, Mya), which are based on the previous report (Stein, et al. 2018). (d-e) The numbers of de novo genes and gene duplicates with different ages across the evolutionary branches.

Leveraging Metapredict, a state-of-the-art deep-learning tool (Emenecker, et al. 2022), our analysis shed light on the intrinsic structural disorder (ISD) of de novo genes (Supplementary table 2). We discovered that 37.57% (68 out of 181) of de novo proteins exhibit complete ISD, characterized by being composed entirely of intrinsically disordered regions (IDRs) (Figure 2a). Notably, this proportion far surpasses the 9.77 % of complete ISD proteins in gene duplicates from age groups br1 to br6 (823 out of 8427). The overall distributions of ISD ratio (the ratio of sequence as IDRs) further showed that de novo genes are strikingly different from gene duplicates in terms of both median value (0.88 vs. 0.31) and distribution peak (0.97 vs. 0.08) (Figure 2b). Interestingly, we found that de novo genes gradually reduce in fractions of IDRs (regions of ISD), suggesting the decay of disorder over evolutionary time (Figure 2c). Specifically, the fractions of IDRs in de novo proteins have decreased by about 40% from the most recent branch (br1) to the oldest one (br6). In addition, de novo genes demonstrated a consistent pattern of higher proportions of IDRs than gene duplicates at all evolutionary stages within ∼1-2 million years (br1-br6), despite a reduced difference between them at the oldest stage br6 (Figure 2c). This pattern suggests that ISD levels in proteins are not stagnant over evolutionary time in rice. Statistically, a significant linear trend emerged: the proportions of IDRs in de novo proteins decrease by about 14% per protein per million years (Figure 2c, *p* = 0.0022, adjusted R^2^ = 0.904). Using the median ISD ratio of gene duplicates (0.31) as a benchmark, and guided by this linear model, de novo proteins would require approximately 4.7 million years to attain the median disorder level observed in gene duplicates.

**Figure 2.**
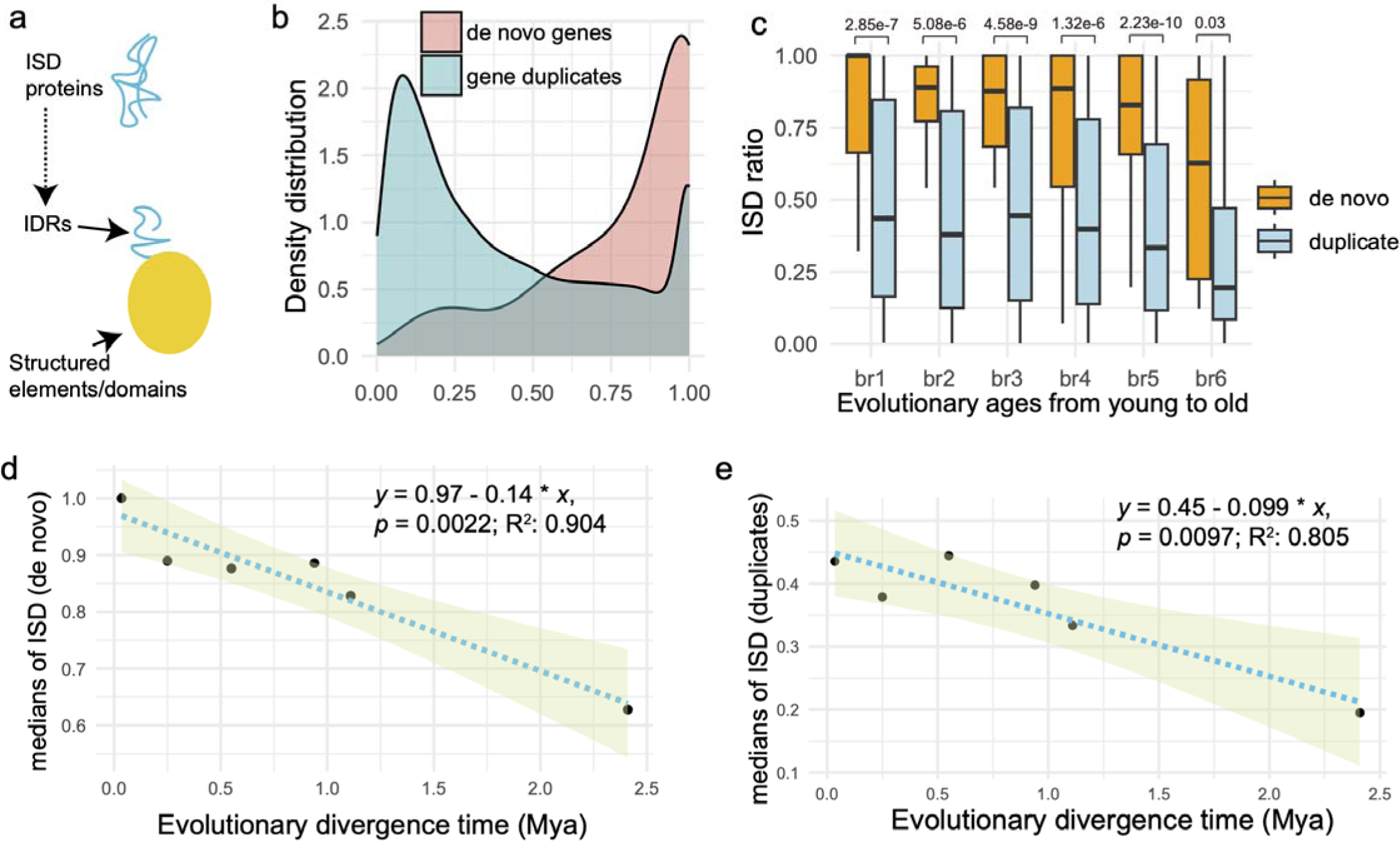
Analysis of intrinsic structural disorder (ISD) in de novo genes and gene duplicates. (a) Illustration of an ISD protein highlighting the intrinsic disordered regions (IDRs). (b) Distribution comparison of IDR fractions in de novo genes versus gene duplicates. (c) Boxplot representation of IDR fractions (also the ISD ratios) in proteins for de novo genes and gene duplicates, categorized by evolutionary age from young to old (x-axis). Differences are assessed using the Wilcoxon test, with the *p*-value indicated above each comparison. (d) A significant linear regression analysis showing the relationship between the median ISD ratios and the evolutionary ages of de novo genes. The 95% confidence interval is represented by the shaded area. (e) Similar linear regression analysis for gene duplicates (br1-br6), with the median ISD fractions plotted against evolutionary ages. The shaded area indicates the 95% confidence interval. The linear regression formula, *p*-value, and adjusted R-squared values are displayed at the top right corner.

For gene duplicates, we found that 19.57% (1818 out of 9289) of proteins encoded by younger duplicates (branches br2-5, ∼1mya) are categorized as ISD proteins (using 100% disorder as the threshold). This rate is 8.4 times higher than that observed in older duplicates from stages br6-9 (2.32%, 570 out of 24620) (Supplementary figure 1). For the *O. sativa Japonica* specific duplicates (br1), we divided the duplicates into two groups: young-parent duplicates and old-parent duplicates, based on the evolutionary epochs from which their parent gene emerged (br2-5 as young parent vs. br6-9 as old parent). Our analysis revealed a significantly higher fraction of ISD proteins in young-parent duplicates compared to old-parent duplicates (58.60%, 53 out of 215 vs. 32.14%, 26 out of 252; Odds ratio 2.38, 95% CI: 1.44 to 3.95, *p* = 0.0007). This finding suggests a “heritage effect,” indicating that gene duplicates may inherit structural properties from their parental genes.

In our comparative analysis of the evolutionary rate of ISD fractions between de novo genes and gene duplicates across branches br1 to br6 (Figures 2d-2e), we uncovered a notable trend. De novo genes exhibit a 4% faster rate of disorder decay per protein per million years than gene duplicates, with respective slopes of 0.14 versus 0.099. This accelerated rate in de novo genes may stem from their absence of the intrinsic heritage effect, which in turn could contribute to their heightened evolvability compared to gene duplicates.

### Rapid evolution of structural elements in de novo proteins

In protein structure, α-helices and β-strands are typically amphipathic and thus can enable the tertiary folding of hydrophilic surfaces and hydrophobic cores (Fersht 1999). The α-helices (and other helices like 3_10_ helices) and β-strands (which form β-sheets) are considered structured due to their specific, stable hydrogen-bonding patterns, while random coil regions lack such regular structure and are more flexible and disordered (Craveur, et al. 2015) (Figure 3a). We conducted a comparative analysis of these structural elements for de novo genes and gene duplicates, focusing on relative proportions of these structural elements within protein sequences over evolutionary time. We predicted protein three-dimensional structures with AlphaFold2 (Supplementary figure 2-7), decoded the structural elements with STRIDE (Heinig and Frishman 2004; Jumper, et al. 2021), and finally measured the lengths and proportions of these structural elements (*P_coil_* for coil, *P_helix_ for* α-helices, *P_310helix_ for* 3_10_ helices, *and P_strand_*for β-strands). Our analysis revealed that median proportion values are highest in unstructured coils (40%-47%) and followed by α-helices (23%-30%), β-strands (13%-15%), and 3_10_ helixes (2.7%-2.8%) for de novo genes and gene duplicates (Supplementary table 3).

**Figure 3.**
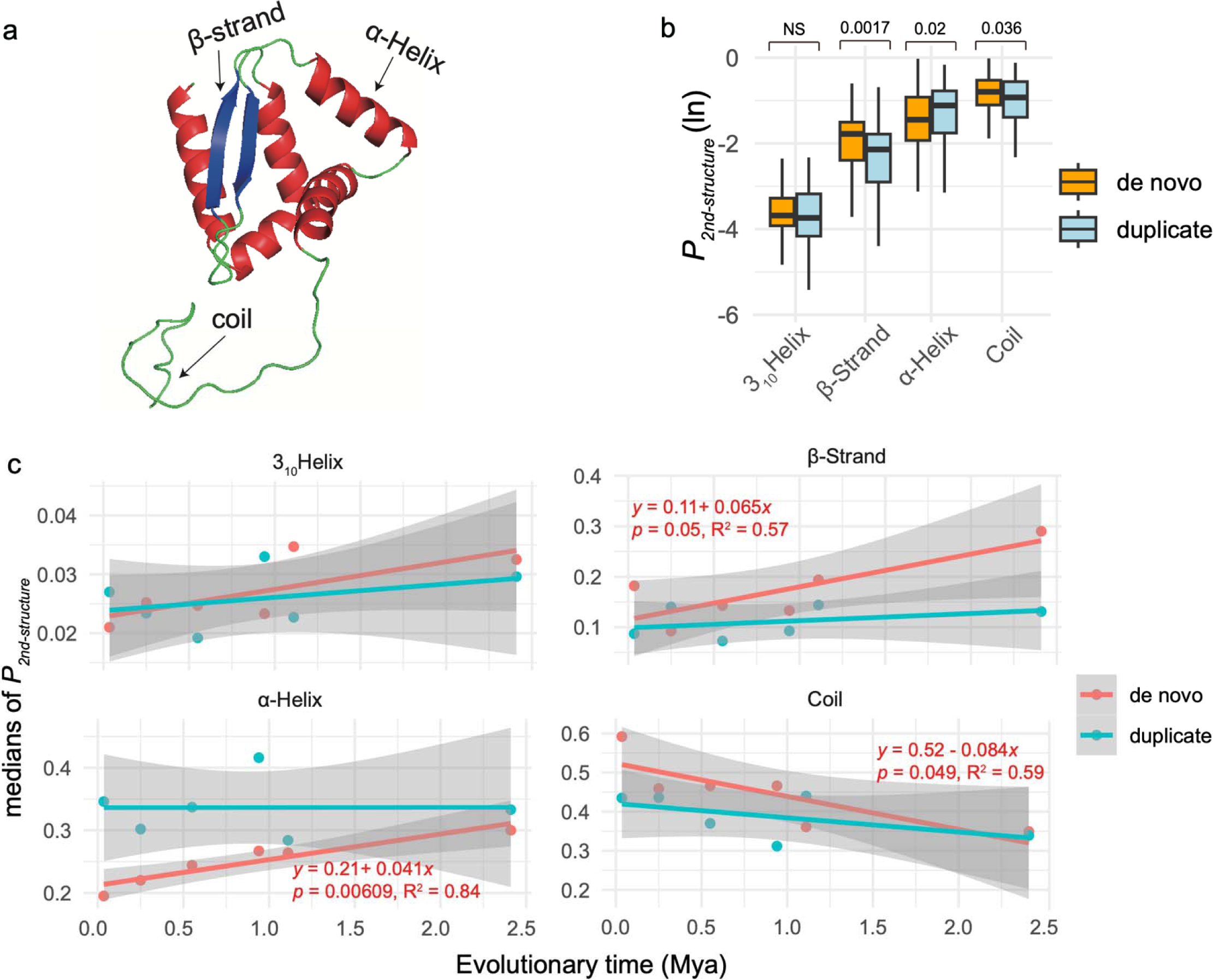
The length proportions of structural elements (noted as *P_2nd-structure_*, transformed using the natural logarithm), including unstructured (coil) and structured segments (3_10_ helix, α-helix, and β-strand) and their correlations with gene ages. (a) An example of basic elements of protein structure. The visualization is based on the ranked_0 result of AlphaFold2 for de novo gene Osjap03g04 70. (b) The distributions and comparisons for length proportions of coil and other structured regions segments (α-helices, 3_10_ helices, and β-strands). The comparisons are based on Wilcox test and *p* values are shown above boxplots. (c) The linear regression of *P_2nd-structure_* for de novo genes against evolutionary time. The linear statistical summaries and formulas are indicated in red for de novo genes. The regression statistics of gene duplicates are not shown due to insignificant *p* values for all elements.

Overall, the *P_coil_*, *P_helix_, and P_strand_* are significantly different between de novo genes than gene duplicates (Figure 3b). In de novo genes, our analysis revealed a strong negative linear correlation between median of *P_coil_* and gene age, alongside significant positive linear correlations between both median of *P_helix_ and P_strand_*and gene age (Figure 3c). These correlations suggest a faster evolutionary rate in the structural elements of de novo genes over time, marked by an increase in novel structures and a decrease in unstructured coil segments. Specifically, α-helix and β-strand grow with rates of 4.1% and 6.5% per protein per million years, respectively, while coil decreases with rate of 8.4% per protein per million years (Figure 3c). In contrast, such correlations are not significant in gene duplicates (Figure 3c). These results indicate a rapid structural evolution in de novo proteins, characterized by a decreasing proportion of disordered or unstructured regions and an increasing proportion of structured regions over evolutionary timescales.

### The properties of amino acids in de novo genes are consistent with the structural changes

The observed patterns for IDRs, random coils, and structured elements (α-helices and β-strands) in de novo proteins necessitate a more comprehensive analysis of amino acid composition to further understand de novo gene evolution. To understand whether the constitutional fractions of some amino acids could be related to gene ages, for each amino acid, we assessed the correlation between median values of fractions and evolutionary ages (Figure 4a). We also compared between de novo genes and gene duplicates (Table 1 and Supplementary figure 8).

**Figure 4.**
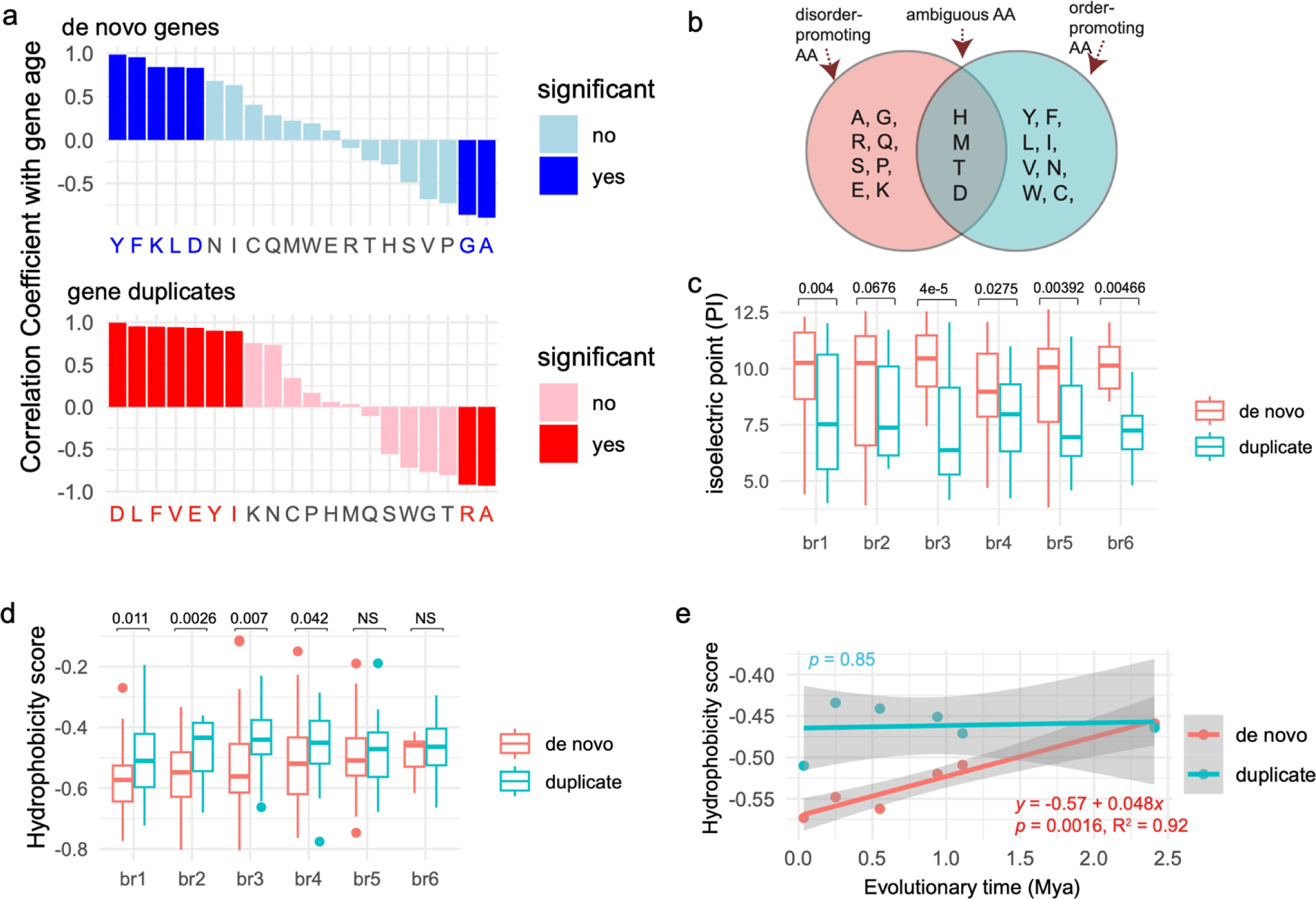
The correlation coefficient between compositions of amino acids and gene ages (Mya). (a) The Pearson correlation coefficients (*r*) between amino acid fractions (medians) and their gene ages (Mya, Supplementary table 4). “Yes” and “No” indicate significant and non-significant *p* values, respectively. (b) The classifications of amino acids (AA): disorder-promoting AA, order-promoting AA, ambiguous AA, based on a previous report (Dunker, et al. 2001). (c) The comparisons of isoelectric point (PI) between duplicates and de novo genes across six branches. (d) The compari ons of hydrophobicity scores between duplicates and de novo genes across six branches. The larger values represent higher hydrophobicity. (e) The linear regression of median hydrophobicity scores against evolutionary times. Statistical summaries are shown near regression lines with *p* values, adjusted R^2^ value, and formula. Comparisons are based on the single-tailed Wilcoxon rank-sum test.

**Table 1.** The comparisons between proteins of de novo genes and duplicated genes. The *p*-values are statistical differences between de novo genes and gene duplicates based on the Wilcox test (significance threshold 0.0025 are adjusted by the multiple test). The field of “Codon Degeneracy” indicates the numbers of codons for the corresponding amino acids.

| Amino Acid | Polarity | Codon Degeneracy | Codons | Charge | Volume | Other Important Properties | Abundance in de novo genes | *p*-value |
| --- | --- | --- | --- | --- | --- | --- | --- | --- |
| G | Nonpolar | 4 | GGT, GGC, GGA, GGG | Neutral | Small | Hydrophobic core | Higher | 3.25E-05 |
| P | Polar | 4 | CCT, CCC, CCA, CCG | Neutral | Small | Proline kinks | Higher | 2.56E-05 |
| R | Polar | 6 | CGT, CGC, CGA, CGG, AGA, AGG | Positive | Large | - | Higher | 1.59E-13 |
| D | Polar | 2 | GAT, GAC | Negative | Small | - | Lower | 2.71E-16 |
| E | Polar | 2 | GAA, GAG | Negative | Medium | - | Lower | 1.23E-03 |
| F | Nonpolar | 2 | TTT, TTC | Neutral | Large | Aromatic ring | Lower | 3.27E-08 |
| I | Nonpolar | 3 | ATT, ATC, ATA | Neutral | Large | Hydrophobic core | Lower | 1.17E-07 |
| L | Nonpolar | 6 | TTA, TTG, CTT, CTC, CTA, CTG | Neutral | Large | Hydrophobic core | Lower | 1.67E-09 |
| N | Polar | 2 | AAT, AAC | Neutral | Small | Amide group | Lower | 7.17E-04 |
| V | Nonpolar | 4 | GTT, GTC, GTA, GTG | Neutral | Medium | Hydrophobic core | Lower | 3.65E-10 |
| Y | Polar | 2 | TAT, TAC | Neutral | Large | Hydroxyl group | Lower | 3.53E-09 |

Among all amino acids, the average fractions Alanine (A) and Glycine (G) exhibited significant negative correlations with ages of de novo genes (Figure 4a, Supplementary table 4). This result suggests that the “disorder-promoting” tendency of Alanine and Glycine could promote the higher ISD and fractions of unstructured coils in young de novo genes (Figure 4b) (Dunker, et al. 2001; Uversky 2013). In gene duplicates, Alanine (A) and Arginine (R) were the two amino acids whose fractions significantly negatively correlated with gene ages (Figure 4a). Interestingly, Arginine (R) has lower disorder propensity than Glycine (G) (Uversky 2013). The difference is consistent with our finding of a higher degree of ISD in de novo genes compared to gene duplicates. Tyrosine (Y), Phenylalanine (F), Lysine (K), and Leucine (L) exhibited significant positive correlations with the ages of de novo genes (Figure 4a and Supplementary table 4), suggesting their roles in the rapid structural evolution of these genes. Notably, 75% (3 out of 4: Y, F, and L) of these amino acids are hydrophobic and order-promoting, with low disorder propensities (Dunker, et al. 2001; Tompa 2002; Uversky 2013). The Lysine (K) is positively charged, which could favor salt bridge to interact with negative charged amino acids or interactions with DNA or RNA (Couso and Patraquim 2017).

Comparative analysis revealed that de novo proteins collectively have significantly higher fractions of Glycine (G), Proline (P) and Arginine (R) than gene duplicates (Supplementary figure 8). These amino acids are characterized by high codon degeneracy and encoded by GC rich codons (Table 1), which is consistent with high GC content in rice de novo genes (Zhang, et al. 2019). In addition, significantly higher fractions of R (Arginine) were found in most evolutionary stages of de novo genes (br1-br5, Supplementary figure 8). Interestingly, de novo proteins have a significantly higher fraction of positively charged amino acid residue R (Arginine) and lower fractions of negative charged Glutamate residue (E) and hydrophobic amino acid residue (F) (Table 1).

### De novo proteins: lighter, positively charged, and increasingly hydrophobic over time

Previous studies conducted on yeast and mammals suggests that new proteins are usually positively charged (Blevins, et al. 2021; Papadopoulos, et al. 2021). Despite these findings, the extent to which this characteristic is pervasive among proteins of varying evolutionary ages remains uncertain. We compared several physiochemical properties, including protein charge, molecular weight, and hydrophobicity, between proteins from de novo genes and gene duplicates across evolutionary stages. By evaluating isoelectric point (PI), we found that de novo proteins exhibit higher positive charges than gene duplicates in all evolutionary age groups (Figure 4c). For molecular weights of proteins (Da), de novo proteins displayed significantly lower values than proteins of gene duplicates in most evolutionary age groups (br2-br6, Supplementary figure 8c).

Interestingly, de novo proteins also showed significantly higher hydrophobicity scores than duplicated proteins at the first four evolutionary stages within 0.94 million years (br1-br4, Figure 4d), and no significant difference was found at br5 (∼1 Mya) and br6 (∼2 Mya) (Figure 4d). Moreover, only in de novo proteins, we detected a significant increasing trend of hydrophobicity score over time with the growth rate of 4.8% per protein per million years (Figure 4e). Due to the dominant role of hydrophobic interactions in driving protein folding, the growth pattern of hydrophobicity strongly supports the faster evolution of folding in de novo proteins, which is also consistent with our findings of secondary structural elements (Figure 3c).

### Protein complex interaction could facilitate the structural evolution of de novo protein

We analyzed the tertiary folding or three-dimensional structure for all de novo genes and a random selection of duplicated genes (30 genes per age group). Based on the per-residue confidence score (pLDDT), a widely used measure of modeling quality in AlphaFold2 (Jumper, et al. 2021), we compared the predicted tertiary folding qualities between de novo genes and gene duplicates (Figure 5a). The median pLDDT scores were consistently higher in gene duplicates than in de novo genes, suggesting a greater confidence in the modeling predictions for the tertiary structures of duplicated proteins (Supplementary figure 9a). This pattern is also consistent with our findings of higher levels of ISD in de novo genes (Figure 2c). We further categorized proteins into three distinct groups based on their folding characteristics, as indicated by pLDDT (Supplementary table 5). We found that 3.43% of de novo genes (6 out of 175) have the high folding quality in at least one element over 10 continuous amino acids (pLDDT > 0.9) and 17.14% of de novo genes (30 out of 175) have elements with confident folding quality (pLDDT > 0.7, Supplementary table 5). Among these predicted well-folded genes, only six genes have two structural elements while most of them have a single structural element (α-helix or β-sheet), consistent with previous observations of limited folding in de novo gene-encoded proteins in other species (Peng and Zhao 2023).

**Figure 5.**
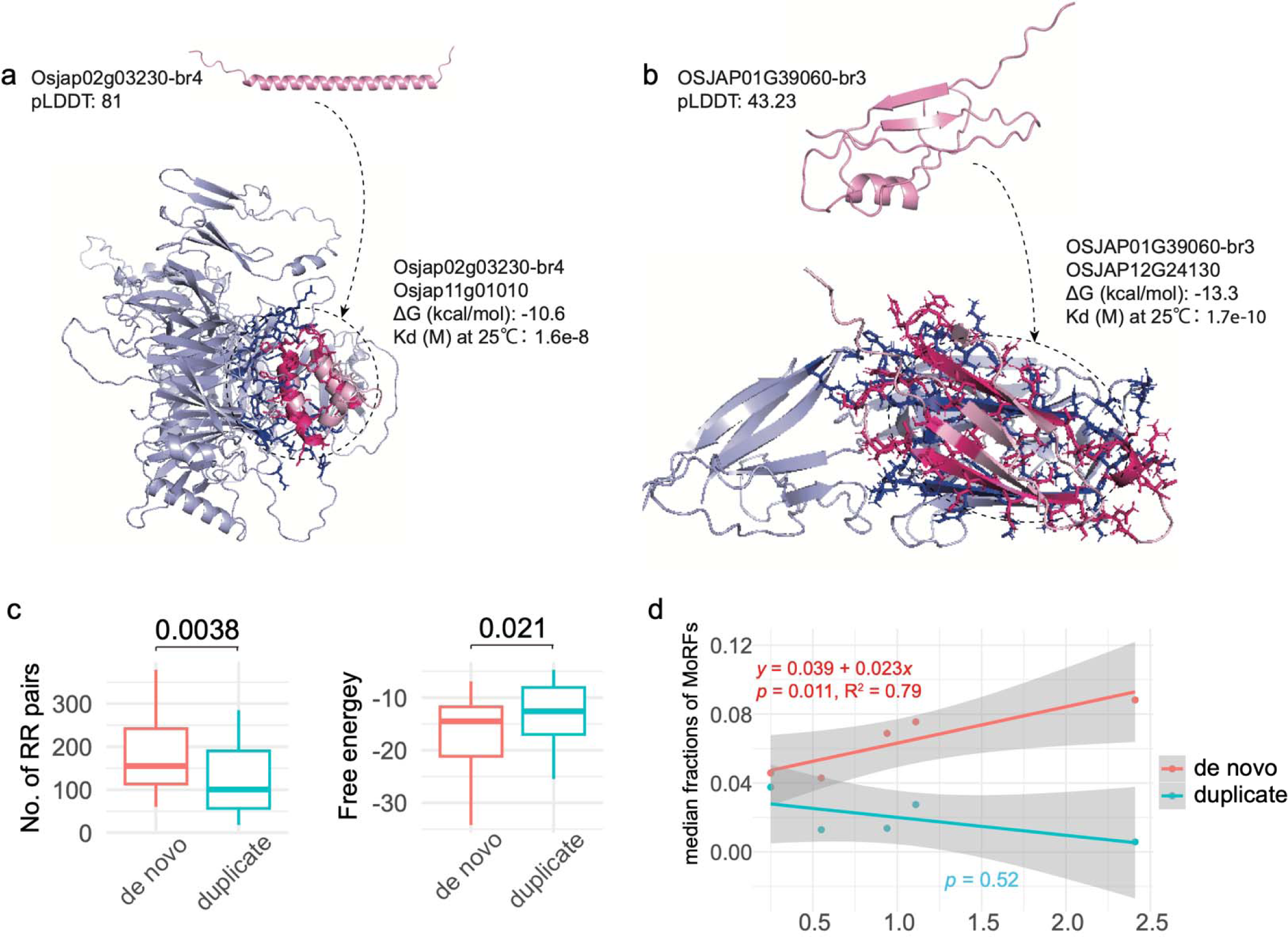
The visualization and statistics of structures for proteins and complexes (a) The 3D structures of Osjap02g03230 and its protein complex. pLDDT indicates average value for all four models, showing a well-folded example. The dotted circle shows the binding state of this de novo protein. (b) The 3D structures of OSJAP01G39060 and its protein complex. pLDDT indicates average value for all four models, representing a not-well-folded example. (c) The comparisons of numbers of residue-residue pairs (RR) and Gibbs free energies (kcal/mol) from results of protein complexes (the model ranked_0) with AlphaFold2-multimer between de novo proteins and duplicates. All comparisons are estimated with the single-tailed Wilcoxon test (*p* values shown above). (d) The regression of linear model between median MoRF fractions and evolutionary years (Mya). The statistical summaries of linear model are listed for the two types of genes (de novo genes and duplicates).

Most proteins function through interactions with other proteins, a process that can induce conformational changes, particularly in disordered proteins (Tsaban, et al. 2022; Zhang, et al. 2013). To explore the likelihood of disorder-to-order transitions during these interactions over time, we assessed the length proportions of molecular recognition features (MoRFs), which are prone to conformational changes during protein-protein contact. Intriguingly, we found that MoRF fractions are consistently higher in proteins from de novo genes than duplicated genes, despite statistical variations for the youngest two age groups (br1 and br2) and older evolutionary ages (br3-br6, Supplementary figure 9b). In de novo genes, we observed a significant linear increase in the median MoRF fractions over evolutionary time, growing at 2.3% per protein per million years (br2-br6, Figure 5b). These findings suggest that the “heritage effect” of duplicated genes could impede the emergence of novel MoRF sequences, while de novo genes could evolve de novo MoRFs for molecular recognition during binding.

Using gene co-expression correlation analysis of RNAseq data (Supplementary table 6) and based on several criteria including disorder levels (ISD proportions < 5%) and high correlation coefficients (> 80%) (See Methods), we identified 30 pairs of potential protein-protein interactions involving de novo proteins (Supplementary table 7). We also used the same criteria and randomly chose 30 pairs of co-expressed gene duplicates for comparison (Supplementary table 7). Using the AlphaFold2-multimer, we tested the possibility of spontaneous de novo protein complexes and potential conformational change upon protein-protein interaction (Bryant, et al. 2022; Evans, et al. 2022; Tsaban, et al. 2022). In one instance, de novo gene Osjap02g03230, which exhibited a highly confident folding structure with a single α-helix, had a predicted conformational change of two α-helices upon binding to its potential protein partner Osjap11g01010, a geranylgeranyl transferase type-2 subunit beta-like protein, with very low free energy (ΔG = -10.6, Figure 5c). The protein complex prediction based on AlphaFold2-multimer indicated a conformational change into a “helix-turn-helix” motif upon binding (Figure 5a). ΔG values are generally in the range of -5 to -10 kcal/mol for biologically relevant interactions (Yugandhar and Gromiha 2014). Thus, the estimate of Gibbs free energy (ΔG) suggests a strong biological relevant binding affinity for this protein complex based on reported cut-off (ΔG around -10) (Nikam, et al. 2023; Yugandhar and Gromiha 2014). Another de novo gene, OSJAP01G39060, showed a stronger binding affinity, as indicated by low ΔG and Kd values (ΔG = -13.3, Figure 5b). Moreover, two more β-strands were observed in this protein complex, supporting the potential structural and conformational change upon binding. These two groups of ΔG and Kd values indicate that the binding processes could be spontaneous and stable for de novo proteins.

Interestingly, using the Gibbs free energy (ΔG) as the indicator of protein-protein binding affinity, we found that de novo proteins have significantly stronger binding affinities with their partners than proteins from gene duplicates (median -16.67 vs. -13.08, single-tailed Wilcoxon rank-sum test, *p* = 0.021, Figure 5c). This observation is also consistent with our finding of significantly more residue-residue (RR) contacts in de novo protein complexes than in those from gene duplicates (median 183 vs. 125, single-tailed Wilcoxon rank-sum test, *p* = 0.0038, Figure 5d). On average, RR pairs were estimated to be 4.71% more in de novo protein complexes than in protein complexes of duplicated proteins (12.22% vs. 7.51%, Supplementary table 7). These results strongly suggest that the disordered and flexible nature of de novo proteins could facilitate strong binding between proteins. Notably, among all 30 pairs of de novo protein interactions studied (Supplementary table 7 and Supplementary figure 10), we revealed only 17% of potential protein complexes (5 out of 30) with ΔG values larger than -10 kca/mol, suggesting that most of de novo genes (83%) can form highly compact and high-affinity complexes with low free energy (Supplementary table 7b). Together, our results suggest that de novo proteins could form stable complexes with biological relevant binding and may even undergo significant conformational changes.

## Discussion

### Emergence of de novo genes: pioneering the new frontiers of evolutionary biology

Over the past decades, the significance of evolutionary new genes has been increasingly recognized, shaping a new paradigm in evolutionary biology (Betrán and Long 2022). Empirical evidence has highlighted the essential and innovative roles of evolutionary new genes in biological systems (Chen, et al. 2013; Heinen, et al. 2009; Long and Langley 1993; Long, et al. 2013; Tautz 2014; Xia, et al. 2021; Xie, et al. 2019a; Xie, et al. 2019b; Zhuang, et al. 2019). One proposed mechanism for the rapid functional diversification of new genes is their low pleiotropic constraints as a competitive advantage compared to older and conserved genes (Chen, et al. 2023a; Hoekstra and Coyne 2007).

Both de novo genes and gene duplicates are important raw materials for evolutionary innovation (Long, et al. 2013), with similar persistence rates in deep evolutionary lineages (Montañés, et al. 2023). As a predominant part of protein-coding genes in genomes, gene duplicates have been modeled to have multiple possible fates, including novel functions (Birchler and Yang 2022; Ohno 1970). However, the possibility of origination and functionalization de novo genes was long dismissed (Jacob 1977; Mayr 1982). Nevertheless, recent studies have provided substantial evidence for de novo gene origination and function (An, et al. 2023; Cai, et al. 2008; Chen, et al. 2023b; Heames, et al. 2020; Qi, et al. 2023; Suenaga, et al. 2014; Zhang, et al. 2019; Zhuang and Cheng 2021).

The structure-function relationship in structural biology suggests that a protein’s primary sequence dictates its tertiary conformation, which in turn influences protein function (Anfinsen and Haber 1961). This underscores the importance of investigating the structural evolution of proteins, particularly in the case of those that arise “from scratch.” With cutting-edge computational tools now available, researchers have begun on detailed case studies to elucidate the foldability and inherent structure of de novo genes (Bornberg-Bauer, et al. 2021; Bungard, et al. 2017; Lange, et al. 2021). Some studies have shown that a fraction of de novo genes can emerge as well-folded and stable entities (Peng and Zhao 2023), while other studies suggest little change over millions of years (Lange, et al. 2021). However, the fundamental question involving the evolutionary pace of structural modifications in de novo genes remains a largely unexplored aspect of evolutionary biology.

### De novo proteins initially exhibit high disorder but rapidly evolve towards structured forms

By comparing our previously identified de novo genes with gene duplicates across well-ordered evolutionary timescales (Zhang, et al. 2019), we observed striking features that the median proportion of intrinsically disordered regions (IDRs) is 88%, indicating disorder as a predominant characteristic for these proteins over a period of 1-2 million years. The structural versatility of IDRs could confer special molecular advantages for de novo proteins, allowing them to adapt to almost every cellular compartment and perform various functions, including transcription, nuclear transport, RNA binding, signaling, and cell division (Holehouse and Kragelund 2023). For instance, numerous RNA binding proteins and transcription factors, which are known to bind nucleic acids and mediate protein-RNA or protein-DNA interactions, contain IDRs (Brodsky, et al. 2020). Another significant example is the IDRs found in eukaryotic histone tails, which undergo post-translational modifications essential for gene expression regulation throughout development (Jiao, et al. 2020). The extensive IDRs we identified in proteins encoded by de novo genes, along with the documented versatility of IDRs in facilitating various interactions and participating in cellular processes, may be pivotal in the evolutionary emergence and functional integration of proteins encoded by de novo genes.

We also found a rapid evolution of their protein structures compared to proteins from gene duplicates within the time frame of 1-2 million years. This rapid evolution is characterized by a decrease in the proportion of unstructured regions (random coils) and an increase in structured regions, such as α-helices and β-strands. We also detected signals of molecular recognition features (MoRFs) and their growing pattern over time. It is known that α-helices and β-strands are both hydrophilic and hydrophobic, which could determine the protein’s tertiary folding by orienting hydrophilic surfaces outward and tucking hydrophobic regions inward (Fersht 1999). We found that, despite strikingly higher proportions of IDRs for de novo proteins, the disorder decay rate is at 14% per protein per million years, which is faster than that in gene duplicates with 9.9% per protein per million years. At this rate, de novo proteins could achieve a structural order comparable to the median levels observed in gene duplicates over a period of around 4.7 million years.

We further observed distinct evolutionary patterns in the basic elements of protein folding. Specifically, we estimated a decrease in random coils at a rate of 8.4% per protein per million years, which suggests a reduction in less structured regions where weaker interactions like Van der Waals forces are predominant. Conversely, there was an increase in α-helices and β-strands at rates of 4.1% and 6.5% per million years, respectively. This increase indicates a shift toward more structured and stable configurations, typically stabilized by hydrogen bonding within the protein’s backbone. The growth in α-helices and β-strands suggests an evolutionary trend towards more hydrogen bond-rich and intricately folded structures, possibly reflecting an increased need for functional specificity and molecular stability. Additionally, we revealed a pattern of increasing hydrophobicity in de novo proteins at 4.8% per protein per million years. This rise in hydrophobicity implies an enhanced role of hydrophobic interactions, which are critical in stabilizing the protein’s tertiary structure by driving the folding process and promoting the interior packing of hydrophobic side chains. This evolving hydrophobic character could be a response to the need for more compact and stable protein structures in diverse cellular environments.

In contrast, we did not find significant evolutionary trends for these metrics in gene duplicates over time, suggesting a more conservative structural evolution compared to de novo proteins. Considering the major role of hydrophobicity in promoting the folding process, our results strongly reveal that de novo proteins could evolve novel tertiary structures at a faster rate than duplicated proteins could. The rapid structural evolution in de novo proteins may be attributed to the combined effects of enhanced hydrophobic interactions, increased hydrogen bonding, and a shift towards more defined secondary structures, reflecting an adaptive advantage in evolving new protein functions and interactions.

### Multiple features of de novo proteins could promote the formation of protein complex

Our analyses indicated several unique physiochemical features of de novo proteins compared to proteins of gene duplicates, which could promote the interactions between de novo proteins and other proteins. Although previous findings in other species have revealed significantly higher positive charges in de novo proteins than other genes (Blevins, et al. 2021; Papadopoulos, et al. 2021), it was unknown whether that is general for all evolutionary ages. Our analyses revealed the general patterns of higher positive charges for de novo proteins than duplicated ones in all evolutionary age groups within ∼2 million years. We also revealed the generally smaller weights of de novo proteins than proteins of gene duplicates. Proteins with greater opposite charges could promote stable binding to form complexes (Hazra and Levy 2022). Thus, the “tiny and attractive” features in terms of weight and charge may suggest a “faster-binding” scenario for de novo proteins, where the nascent de novo proteins could have relatively higher diffusion speed to be attracted to the negative charged compartments or larger molecules (Figure 6a). Generally, larger negative charged proteins tend to offer greater collision cross-sections for interactions, while smaller positively charged proteins, with their faster diffusion, are more prone to molecular collisions (Morris, et al. 2022; Xu, et al. 2013). Therefore, our results suggest that de novo proteins, exhibiting generally positive charge and smaller size, may have a higher diffusion potential, increasing their likelihood of interacting with larger, negative charged proteins or cellular structures. The unique physiochemical properties of these nascent proteins could facilitate the formation of novel and diverse protein complexes.

**Figure 6.**
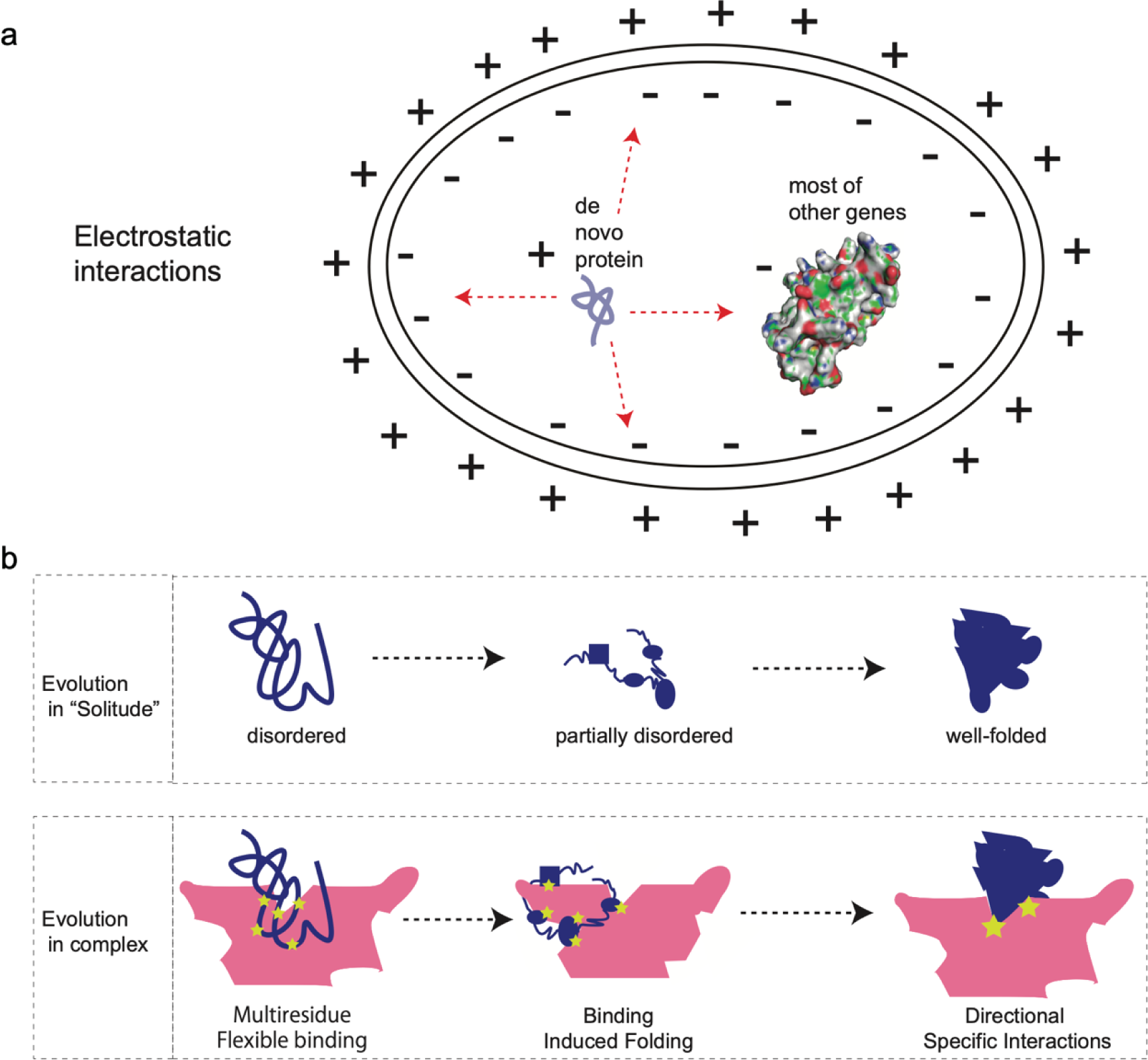
The schematic illustration for molecular diffusion and structural evolution of de novo proteins. (a) The schematic molecular diffusion and movement showing differences in diffusion speed based on protein charges and molecular weight differences between de novo genes and duplicates (also see Supplementary figure 8c for molecular weight differences). The “+” indicates the general positive charges in de novo proteins and outside of the cell membrane. The “-” indicates the more negatively charged proteins from duplicates and the inner side of the cell membrane. The size difference indicates the general pattern of significantly less molecular weight in de novo genes than in gene duplicates. (b) Two models of protein folding evolution for de novo protein: the evolution in “solitude” model (EIS) and the evolution in complex model (EIC).

Our three-dimensional analyses on de novo proteins and complexes revealed contrasting patterns between the isolated protein structure and protein complex. Consistent with the expectation based on high levels of ISD in de novo proteins and findings in other species (Peng and Zhao 2023), we found that the tertiary structures of de novo genes in isolation are simple with limited number of structural elements and not well-folded in general. Only a tiny percent (3.43%) of de novo protein had confidently modeled folding structures based on Alphafold2. This general feature could reflect the nature of disorder propensities of de novo proteins. Surprisingly, however, AlphaFold2-Multimer analyses suggested that most de novo protein complexes (83%) have high binding affinities (Gibbs free energy < -10), despite the disordered nature of de novo proteins in isolation. The binding process also demonstrated the potential conformational changes and enhanced residue-residue contacts for de novo proteins upon interaction. Probably constrained by the rigid bodies of well-folded conserved proteins, interfaces of protein-protein interaction are generally controlled by a small and complementary set of contact residues that maintains most of the binding affinity (Clackson and Wells 1995). Different from the interactions between rigid bodies, the soft bodies of de novo proteins could allow for more contact residues and wider surfaces, thereby leading to high binding affinities between proteins.

Therefore, our results further suggest two complementary models for structural evolution of de novo proteins: the evolution in solitude (EIS) and the evolution in complex (EIC) (Figure 6b). The EIS model emphasizes the intuitive and isolated way of structural evolution step by step over evolutionary time from disordered to partially disordered and then to well-folded. Some distinguished features of de novo proteins, including high positive charges (Figure 4c), small molecular weights (Supplementary figure 8c), more residue-residue contacts in complexes (Figure 5c left), lower free energy in complexes (Figure 5c right), and widespread strong binding for most of de novo proteins (>83%), allow for the second model EIC that emphasizes the role of protein complex comprised of de novo protein and well-folded protein in inducing the evolution of folding domains. The EIC model is also consistent with the previous findings that folding is not necessary for binding (Chebaro, et al. 2015) and network hub proteins tend to be disordered (Haynes, et al. 2006; Midic, et al. 2009). In EIC model, the formation of de novo protein complex could be instant and unspecific after protein emergence, much earlier than the formation of well-folded protein structure in isolation. The EIC model suggests that the tertiary structure evolution of de novo proteins could go through steps from the multi-residue binding (Figure 5c), the binding-induced folding (Figure 5a-5b), and to potentially directional specific binding. The binding-induced folding might be a key mechanism facilitating the rapid decrease in disorder within de novo proteins, presenting an intriguing area for future research.

Overall, our study challenges the traditional view of slow protein evolution by demonstrating that de novo genes can evolve rapidly in structural elements within a relatively short evolutionary timeframe. Although gene duplicates represent over 70% of protein-coding genes, de novo genes in general have faster evolutionary rate in structural changes which highlight the importance of de novo gene emergence as a distinguished source of genetic innovation in organisms. The faster-binding of de novo genes prior to their well-folded structures could be one of mechanisms through which de novo genes are fixed in population, evolve rapidly to acquire new functions, and integrate into existing biological networks by protein-protein interactions. Despite these intriguing patterns, all analyses of this study are based on the *in-silico* estimation which could pose some potential limitations. Future research in this area could provide further insights into the mechanisms driving the rapid evolution of de novo genes and their impacts on the evolution of complex biological systems.

## Conclusion

Our research indicates distinct patterns of rapid structural transformation in de novo genes over a relatively brief evolutionary timeframe of 1-2 million years. Additionally, we estimate that de novo proteins require no longer than five million years to attain an intrinsic structural order comparable to that observed in gene duplicates. Exceptional characteristics of de novo genes, such as their low molecular weights, positive net charges, and strong binding affinities, and more residue-residue contacts, likely drive their efficient diffusion and interactions with other molecules, which are essential for their evolution of biological functions. Hence, our findings highlight the unique mechanisms by which these continuously emerging de novo proteins could escape from a prolonged period of solitary existence in evolutionary history.

## Methods

The de novo gene list and origination branches (ages) were retrieved from a previous study (Zhang, et al. 2019), which was based on the synteny alignment between focal species *O. sativa japonica* (br1) and outgroup species. Based on the Oryza phylogenetic tree, the 11 species were assigned to six age groups for de novo genes: *O. rufipogon* (br2), *O. sativa* subspecies *indica* and *O. nivara* (br3), *O. glaberrima* and *O. barthii* (br4), *O. glumaepatula* (br5), and *O. meridionalis* (br6). The divergence time was based on the previous report (Stein, et al. 2018). The gene duplicates were identified based on BLASTP comparison of genome-wide protein sequences (-evalue 0.001 -seg yes). The gene ages for these genes were determined with a two-step synteny-based method: 1) the reciprocal best orthologous genes were exhaustively searched between focal species and outgroup species; 2) the gene synteny blocks were then constructed based on a criterion of no more than 5 genes within the range of reciprocal best pairs. Due to the higher number of duplicated genes, the groups were further extended into another 3 branch groups, which are *O. punctata* (br7), *O. brachyantha* (br8), and *L. perrieri* (br9).

The genome reference and gene annotations (v66) were downloaded from the Gramene database (http://ftp.gramene.org/oge/release-current/) (Gupta, et al. 2016). All RNAseq short-reads data sequenced with the Illumina platform for Oryza sativa were downloaded from NCBI SRA database (∼400GB bases, 08-25-2023, Supplementary table 6). We filtered the samples with fastp (Chen, et al. 2018) and mapped cleaned reads to the genome reference using STAR v2.7.0a (Dobin, et al. 2013). The expression level for all genes and isoforms were measured with RSEM (Li and Dewey 2011). Since co-expression analysis often involves the relationships between genes across multiple samples, transcripts per million (TPM) was chosen to measure expression because it’s commonly used for inter-sample comparisons. The gene co-expression was analyzed with the Pearson test. We defined the co-expression gene partners (CGP) as the top 30 co-expressed genes with significant interaction signals for each de novo genes (*p* < 10^-5^). We also randomly picked up 200 duplicated genes for comparison.

The intrinsic structural disorder (ISD) of protein-coding genes for rice genome (http://ftp.gramene.org/oge/release-current/) (Gupta, et al. 2016) was analyzed with metapredict (v2.3), a deep-learning based consensus predictor (Emenecker, et al. 2021). ISD proteins were defined as proteins with over 100% of residues under disordered states (threshold 1). The ISD level or proportion was evaluated with the fraction of ISD segment out of the full length of a protein. We performed a linear regression analysis on the median ISD levels of proteins across different evolutionary stages, using the ’lm’ function in the R platform, to assess their relationship with evolutionary time.

The three-dimensional structures of de novo proteins were firstly estimated with AlphaFold2 with default parameters and the structural elements were extracted with STRIDE (Heinig and Frishman 2004; Jumper, et al. 2021). For gene duplicates, we randomly picked 30 genes from each branch. To elucidate the evolutionary dynamics of protein structure, we quantified the proportion of unstructured (random coil) and structured (α-helices and β-strands) regions in both de novo genes and gene duplicates (*P_2nd-structure_*). These proportions are defined by the equations:

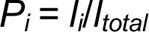

Where *i* represents coil, α-helix, 3_10_ helix, or β-strand, the *l_i_*is the cumulative **l**ength of each element *i*, and *l_total_*denotes the total protein **l**ength. The median values for *P_i_*were used to conduct linear regression against the evolutionary time with R platform.

Molecular recognition features, commonly known as MoRFs, are prevalent components found within disordered regions of proteins, which could transform from a disordered to an ordered state when they bind to their respective protein partners. We predicted the MoRFs using fMoRFpred and compared their proportions between gene duplicates and de novo genes (Yan, et al. 2016). The online tool of ipc2 was used to evaluate isoelectric point (PI) and molecular weights (Da) for all de novo genes and 200 duplicated genes randomly selected (Kozlowski 2021). The hydrophobicity scores were estimated with the previous reported method (Wilson, et al. 2017).

We further classified protein 3D-structures based on Alphafold2 into three groups. The high-confidence potential folding was defined as at least one element over 10 amino acids with pLDDT > 0.9. The medium-confidence folding was defined as at least one element over 10 amino acids with pLDDT > 0.7. Others are defined as low-confidence quality folding. To understand whether the folding conformation could be changed upon protein binding, we chose both highly-confidence folding and low-confidence folding genes and their potential protein partners to conduct protein-protein docking analysis with AlphaFold2-Multimer (Evans, et al. 2022). The protein partners were chosen based on the criteria: 1) low percentage of disordered regions (< 5%); 2) highly correlated expression pattern (co-expression correlation coefficient > 0.8); 3) partner sequence over 200 amino acids and under 500 amino acids. 4) partner as a relatively old gene (br6-br9). The binding free energy and the dissociation constant were estimated with PRODIGY (Vangone and Bonvin 2015; Xue, et al. 2016). The spontaneity and stability of the binding process for protein-protein interactions was evaluated with the change in Gibbs free energy (ΔG) and the dissociation constant (Kd). The cutoff ΔG = −10 kcal/mol (Kd of 10^−8^ M) was used to indicate high affinity (Nikam, et al. 2023; Yugandhar and Gromiha 2014). Generally, a lower Kd value (< 1) and a very negative ΔG indicate a more stable and tightly bound complex (Supplementary figure 6b). Because the residue-residue (RR) pairs or contacts could occur between a residue in one protein and multiple residues of its partner, we counted RR as both raw numbers and nonredundant ratios. The raw numbers were based on number of total RR pairs estimated with the tool PRODIGY, while the nonredundant ratios were estimated by focusing on unique pairs and adjusted with total protein length of complex.

## Acknowledgments

The study was supported by Guggenheim Fellowship of Manyuan Long.

## Supplementary figures and legends

**Supplementary figure 1.**
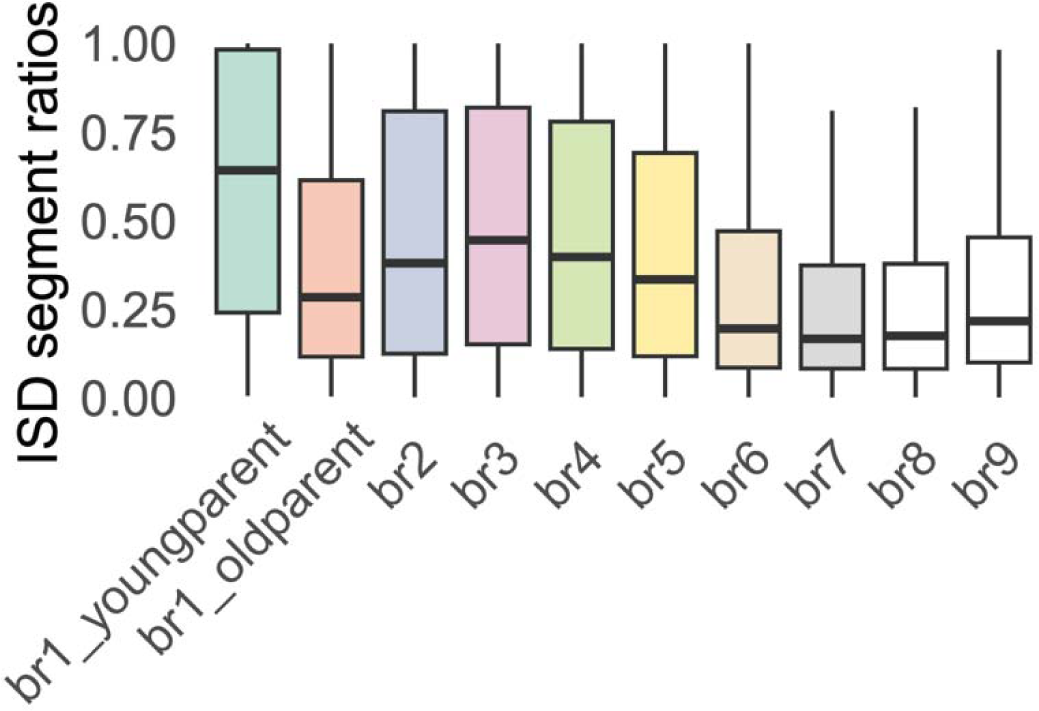
The fractions of intrinsic structure disordered regions for duplicated genes across evolutionary ages from young to old. The br1_youngparent and br1_oldparent indicate young gene duplicates at br1 with parental genes from br2-5 and br6-9, respectively.

**Supplementary figure 2.**
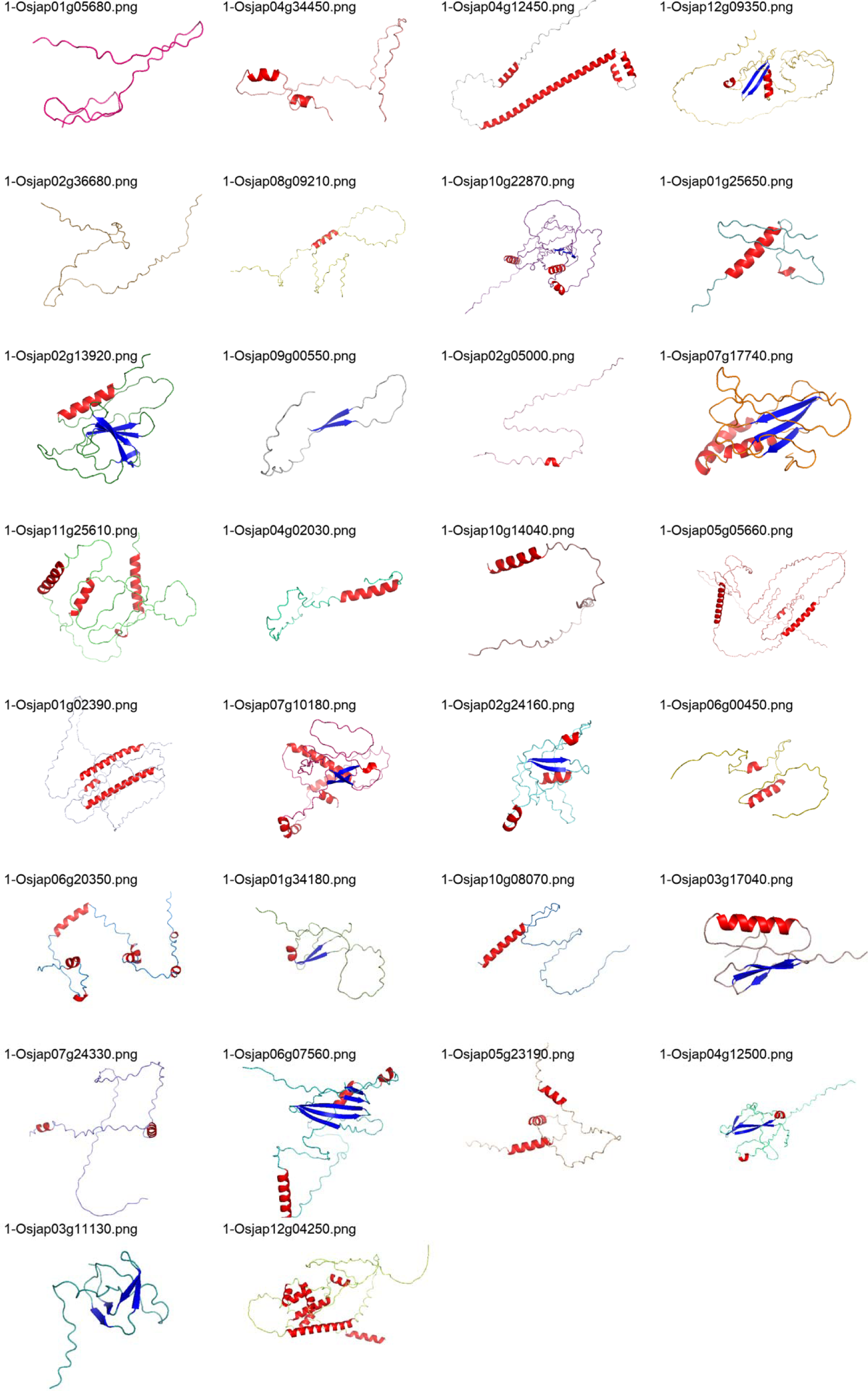
The protein tertiary structure of de novo genes at br1 predicted with AlphaFold2 (ranked_0). The different colors show the predicted different elements (random coil, α-helices, and β-strands). The folding qualities and categories were listed in Supplementary table 5.

**Supplementary figure 3.**
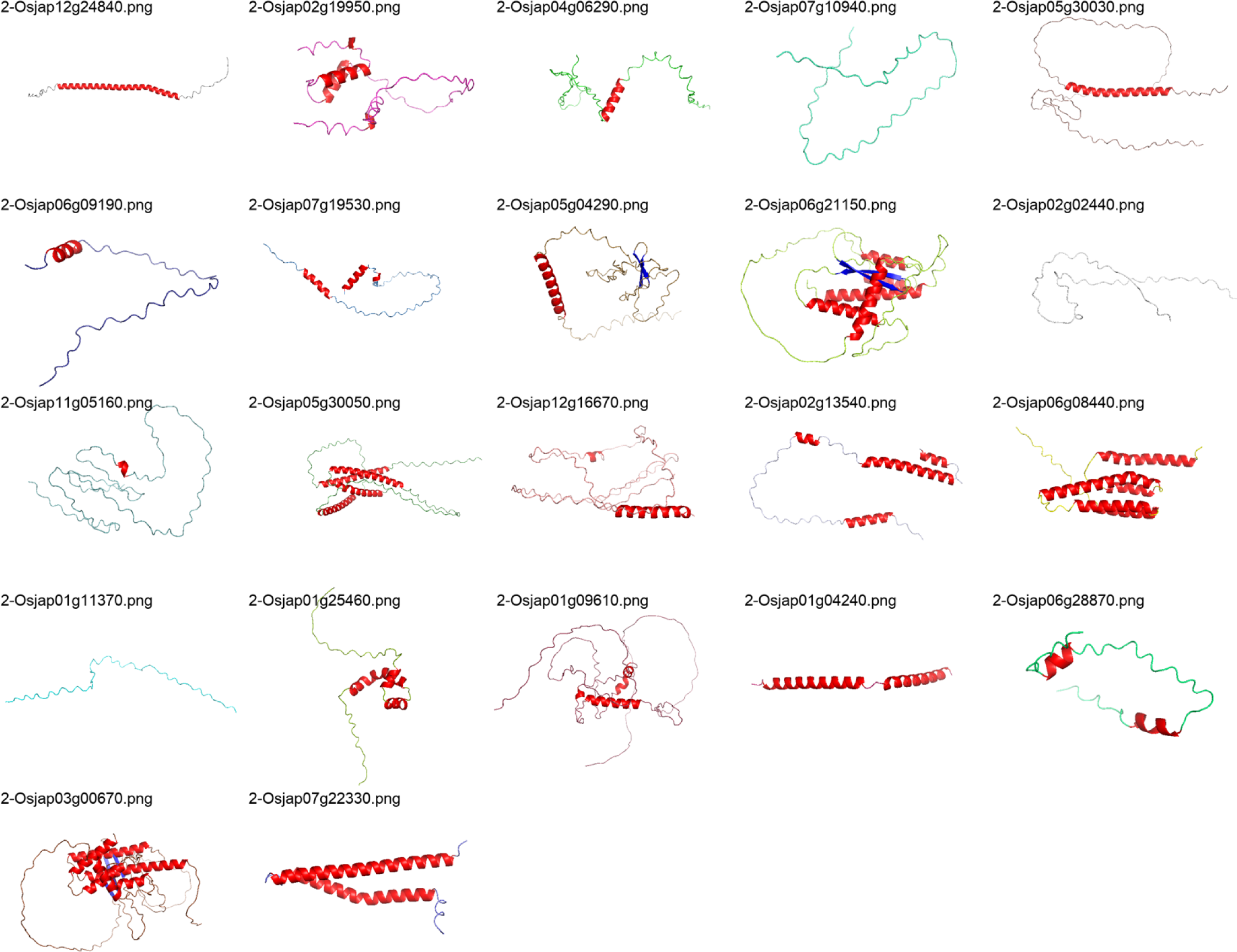
The protein tertiary structure of de novo genes at br2 predicted with AlphaFold2 (ranked_0). The different colors show the predicted different elements (random coil, α-helices, and β-strands). The folding qualities and categories were listed in Supplementary table 5.

**Supplementary figure 4.**
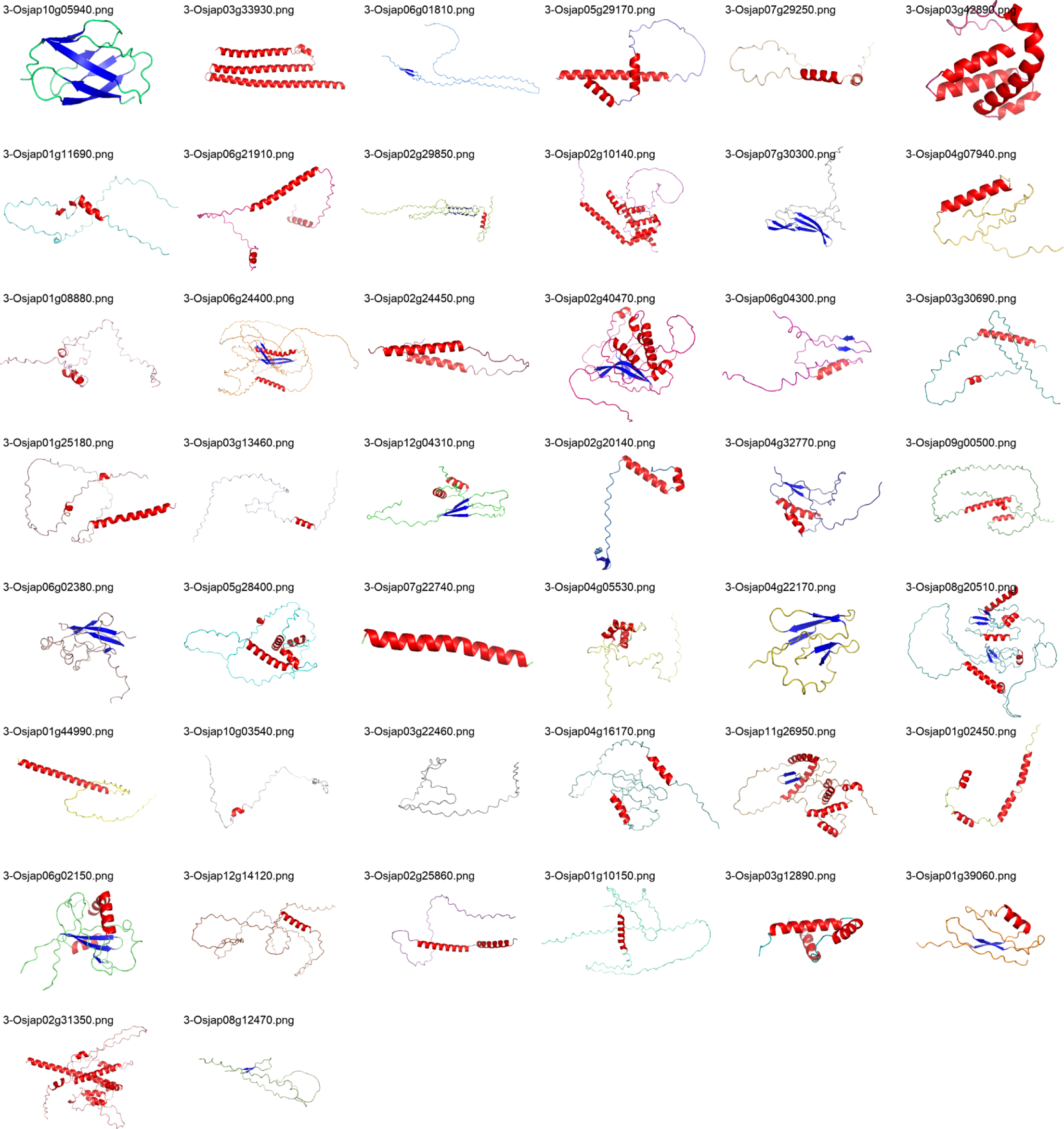
The protein tertiary structure of de novo genes at br3 predicted with AlphaFold2 (ranked_0). The different colors show the predicted different elements (random coil, α-helices, and β-strands). The folding qualities and categories were listed in Supplementary table 5.

**Supplementary figure 5.**
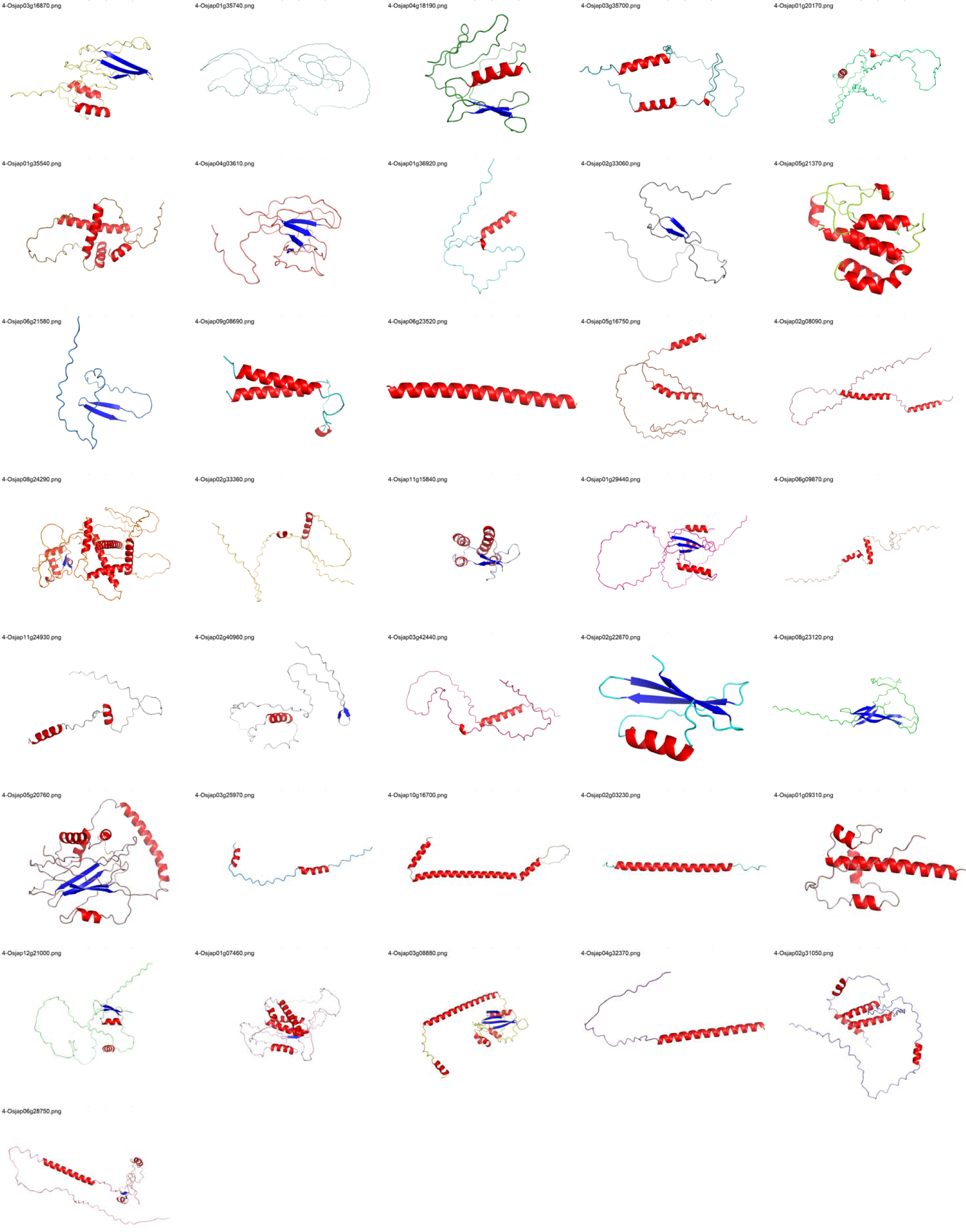
The protein tertiary structure of de novo genes at br4 predicted with AlphaFold2 (ranked_0). The different colors show the predicted different elements (random coil, α-helices, and β-strands). The folding qualities and categories were listed in Supplementary table 5.

**Supplementary figure 6.**
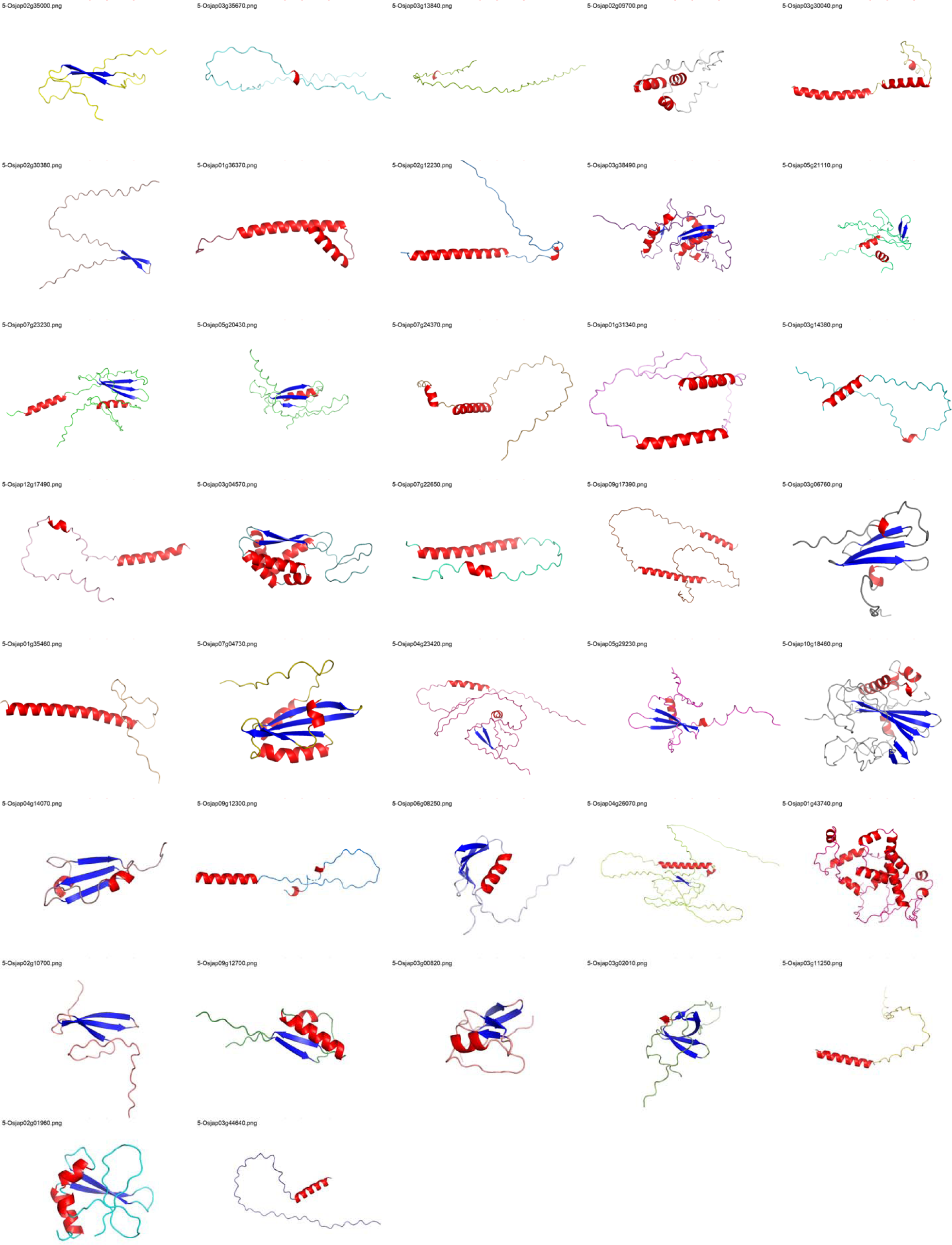
The protein tertiary structure of de novo genes at br5 predicted with AlphaFold2 (ranked_0). The different colors show the predicted different elements (random coil, α-helices, and β-strands). The folding qualities and categories were listed in Supplementary table 5.

**Supplementary figure 7.**
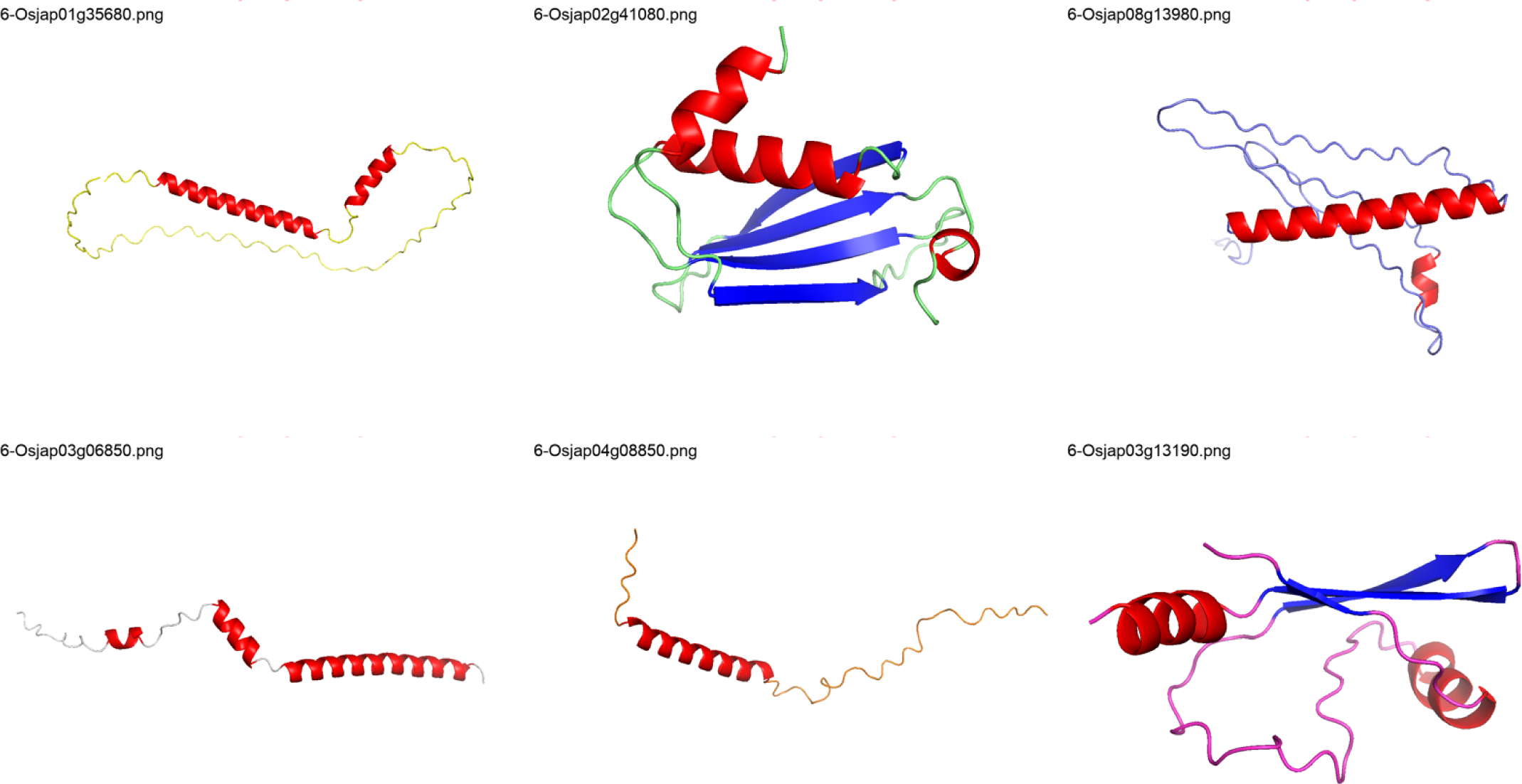
The protein tertiary structure of de novo genes at br6 predicted with AlphaFold2 (ranked_0). The different colors show the predicted different elements (random coil, α-helices, and β-strands). The folding qualities and categories were listed in Supplementary table 5.

**Supplementary Figure 8.**
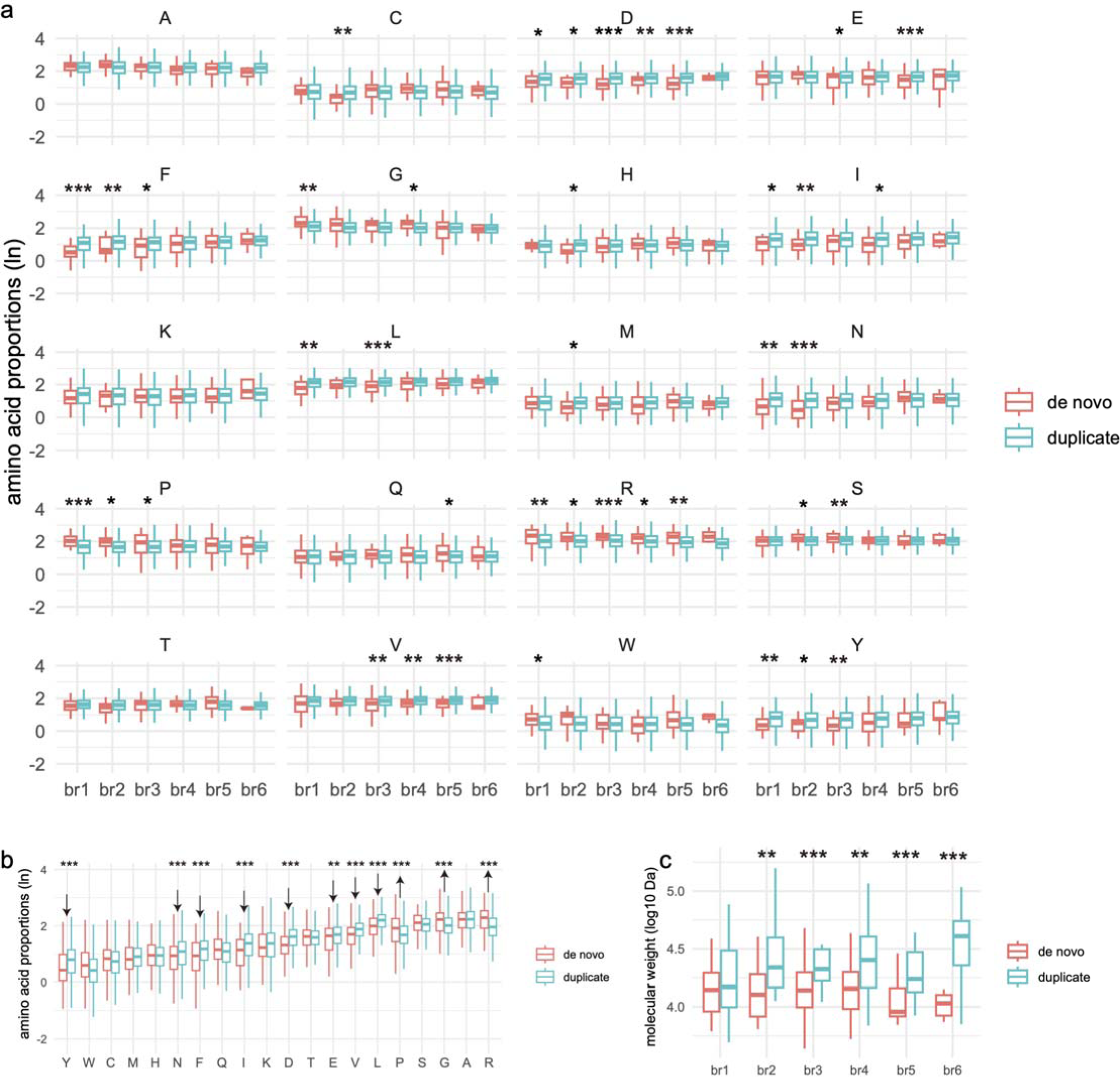
Comparisons of amino acid compositions between de novo genes and gene duplicates across evolutionary branches. (a) Comparisons of the compositions (expressed natural logarithm of percentages) between de novo genes and gene duplicates for different as the amino acids within various branches. (b) Overall comparisons of compositions (natural logarithm of percentages) between de novo genes and gene duplicates for different amino acids, without age differentiation. Arrows pointing upward indicate significantly higher medians in de novo genes, and vice versa. (c) The comparisons of molecular weight (logarithm) between de novo genes and gene duplicates for genes of different ages. Note: All comparisons are based on the Wilcoxon test and only the significant pairs are shown (’’*”, *p* < 0.05; ’’**”, *p* < 0.01; ’’***”, *p* < 0.001).

**Supplementary Figure 9.**
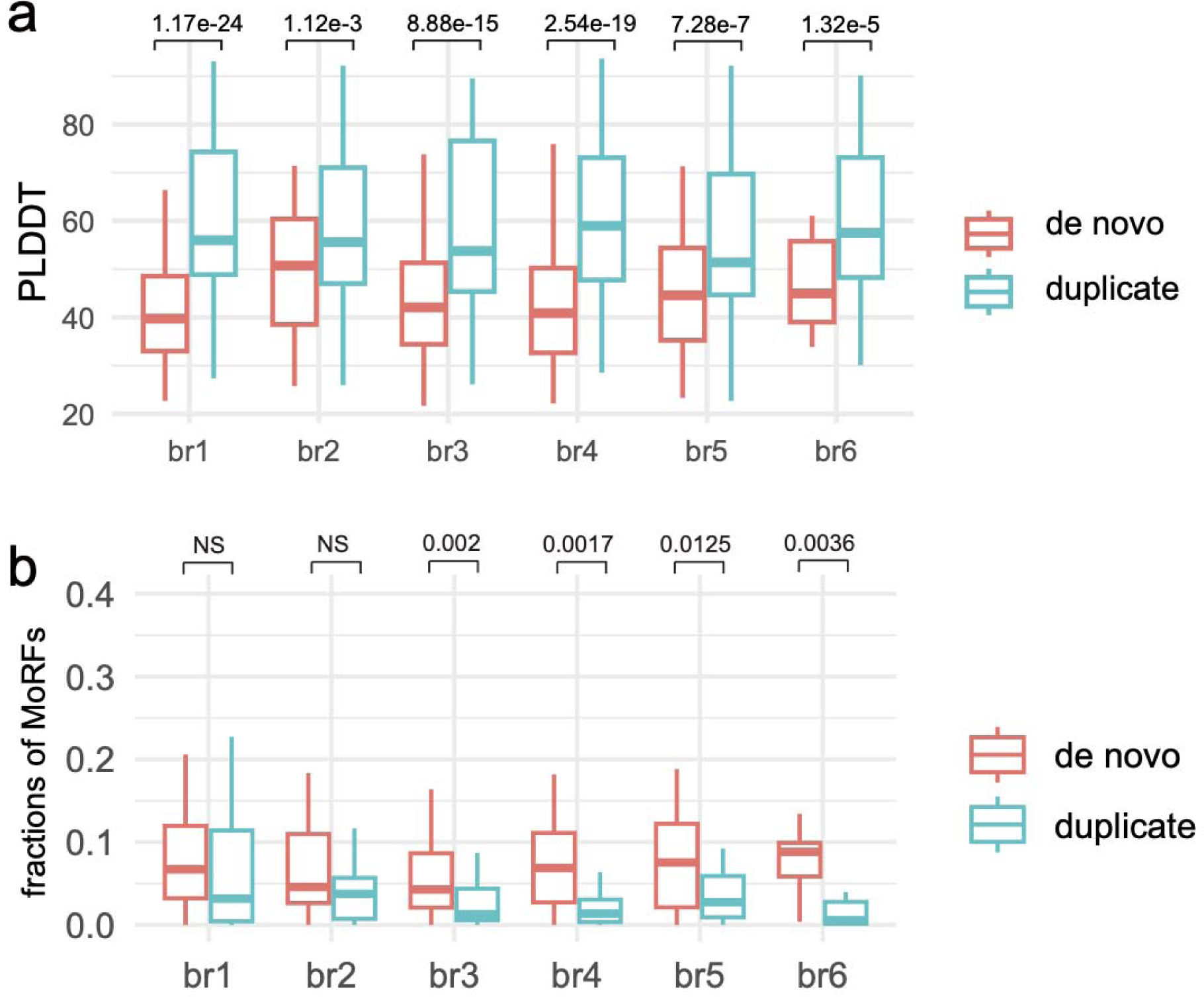
Comparisons of pLDDT scores and Gibbs free energies between de novo genes and gene duplicates. (a) pLDDT score comparisons between de novo proteins and duplicates for folding predictions of isolated proteins with AlphaFold2 across evolutionary branches (ranked_0 to ranked_4). (b) The comparisons of proportions of MoRFs between de novo proteins and duplicates with the Wilcox test (*p* values are shown above).

**Supplementary Figure 10.**
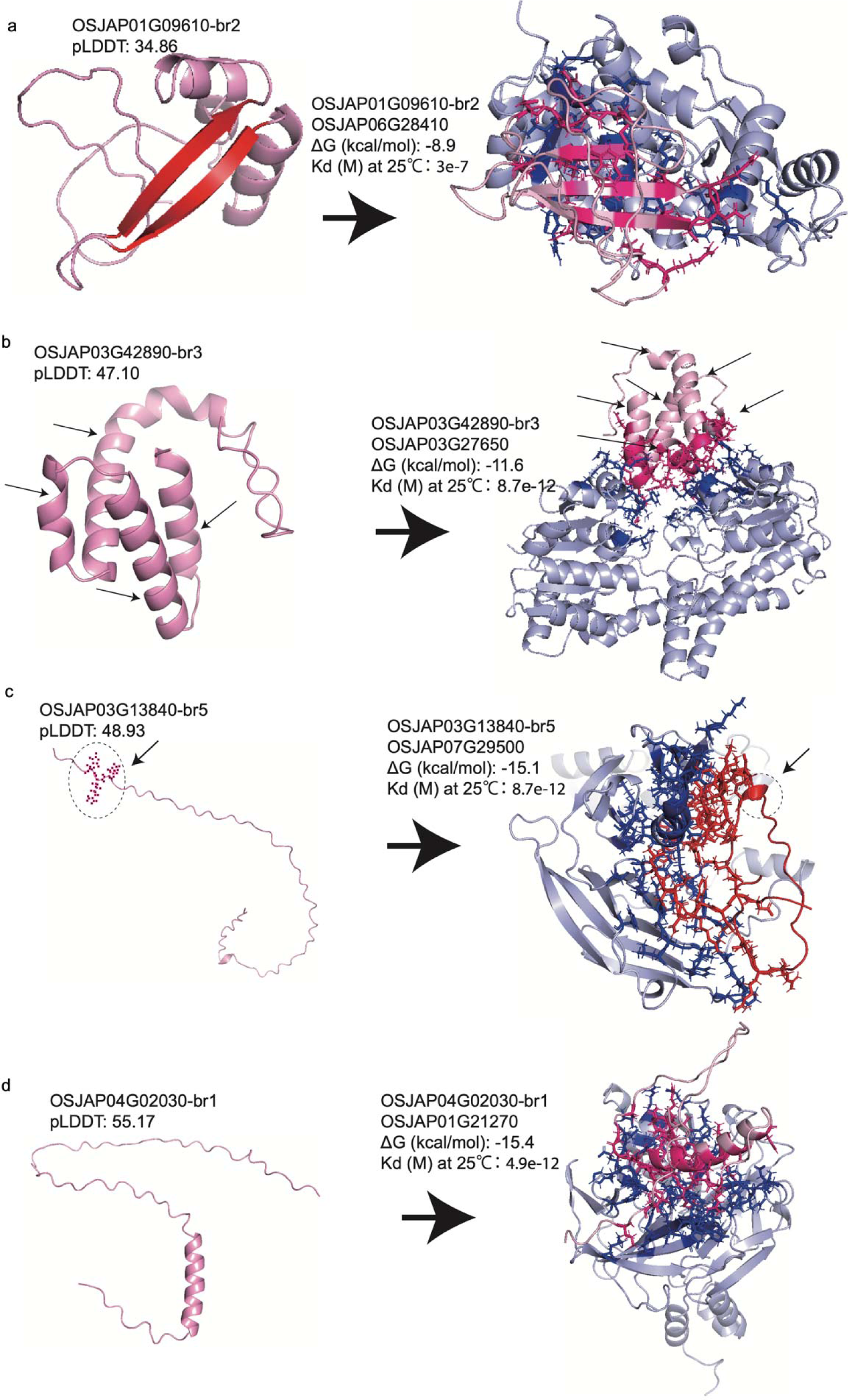
The four examples of 3D structures of de novo genes and their protein complex with binding affinities (Supplementary table 7). pLDDT indicates modeling quality for the model of ranked_0 from AlphaFold2. (a) A new β-strand appears upon binding relative to protein in isolation. (b) Two more α-helices appear in protein complex (shown with arrows). (c) The random coil segment changes into an α-helix in protein complex (shown with arrows). (d) No visible change from single protein to the complex for de novo protein.

